# AGC: Compact representation of assembled genomes

**DOI:** 10.1101/2022.04.07.487441

**Authors:** Sebastian Deorowicz, Agnieszka Danek, Heng Li

**Affiliations:** Faculty of Automatic Control, Electronics and Computer Science, Silesian University of Technology, Akademicka 16, 44-100 Gliwice, Poland; Department of Data Sciences, Dana-Farber Cancer Institute, Boston, MA, USA; Department of Biomedical Informatics, Harvard Medical School, Boston, MA, USA

## Abstract

High-quality sequence assembly is the ultimate representation of complete genetic information of an individual. Several ongoing pangenome projects are producing collections of high-quality assemblies of various species. Here, we show how to represent the sequenced genomes in 2–3 orders of magnitude smaller space, allowing easy and fast extraction of any contig or its part.

---

Rapidly evolving long-read sequencing technologies such as Pacific Biosciences and Oxford Nanopore have enabled routine haplotype-resolved assembly of haploid and diploid genomes [12, 4]. We have started to sequence and de novo assemble collections of samples from the same species [7, 11, 2, 10, 5]. For example, the Human Pangenome Reference Consortium (HPRC) has released 94 haplotype assemblies and plan to produce additional 600 assemblies in the next few years [11]. These haplotype assemblies do not only encode small variants but also represent complex structural variations in segmental duplications and centromeres, empowering the investigation of genetic sequence variations at full scale for the first time. At present, we use generic compression tools, such as gzip, to compress collections of similar genomes. Disregarding high similarity between genomes, these tools can only achieve a 4-fold compression ratio. NAF [9], HRCM [14] and MBGC [6] are the few compression tools that take similarity into account and work with de novo assemblies. However, these tools only aim at reducing transfer and archival costs and are unable to extract individual sequences without decompressing the entire archive. As a result, users have to store uncompressed data for routine analysis. This severely limits their practical applications.

In this paper, we present AGC (Assembled Genomes Compressor), a highly efficient compression method for the collection of assembled genome sequences of the same species. The compressed collection can be easily extended by new samples. AGC offers fast access to the requested contigs or samples without the need to decompress other sequences. The tool is implemented as a command-line application. Access to the data is also possible using C, C++, and Python programming libraries.

The main contribution of the algorithm is the way it represents the assembled genomes and how it supports fast access to the compressed data. Compression is a three-stage process. Initially, a reference genome (provided by the user) is analyzed to find unique *candidate k*-mers (short sequences of length *k*; 31 by default). In the second stage, the reference is analyzed one more time to find *splitters*, which are candidate *k*-mers distant (in the contigs) from each other by approximately *segment size* (60 kbp by default). Proper compression is performed at the third stage, which is executed separately for each added genome. Here, AGC uses splitters to divide each contig into segments. Then, the segments are collected in groups using pairs of terminating splitters to have in the same group segments that are likely highly similar to each other. The actual implementation is more complicated to allow AGC to handle segments with only one terminating splitter (at contig boundaries) or in a situation in which some splitter is ‘missing’ (e.g., due to some evolutionary event). Some insight into such situations is given in Figure 1 and more details can be found in the Methods section. The first segment of a group serves as a reference. The remaining segments are processed in blocks (default size 50). Each segment is represented as descriptions of similarities and differences with respect to the reference segment. These (usually short) descriptions are concatenated and compressed with a general-purpose zstd compressor. This allows a highly efficient representation of not only the similarities between segments and the reference segment, but also among non-reference segments. In the decompression of a single contig, it suffices to read the information about the groups and blocks containing the segments of the requested contig. Then, AGC decompresses the reference segments and, partially, also the necessary blocks. In general, the larger the block size, the better the compression ratio, but the longer the access time.

**Figure 1:**
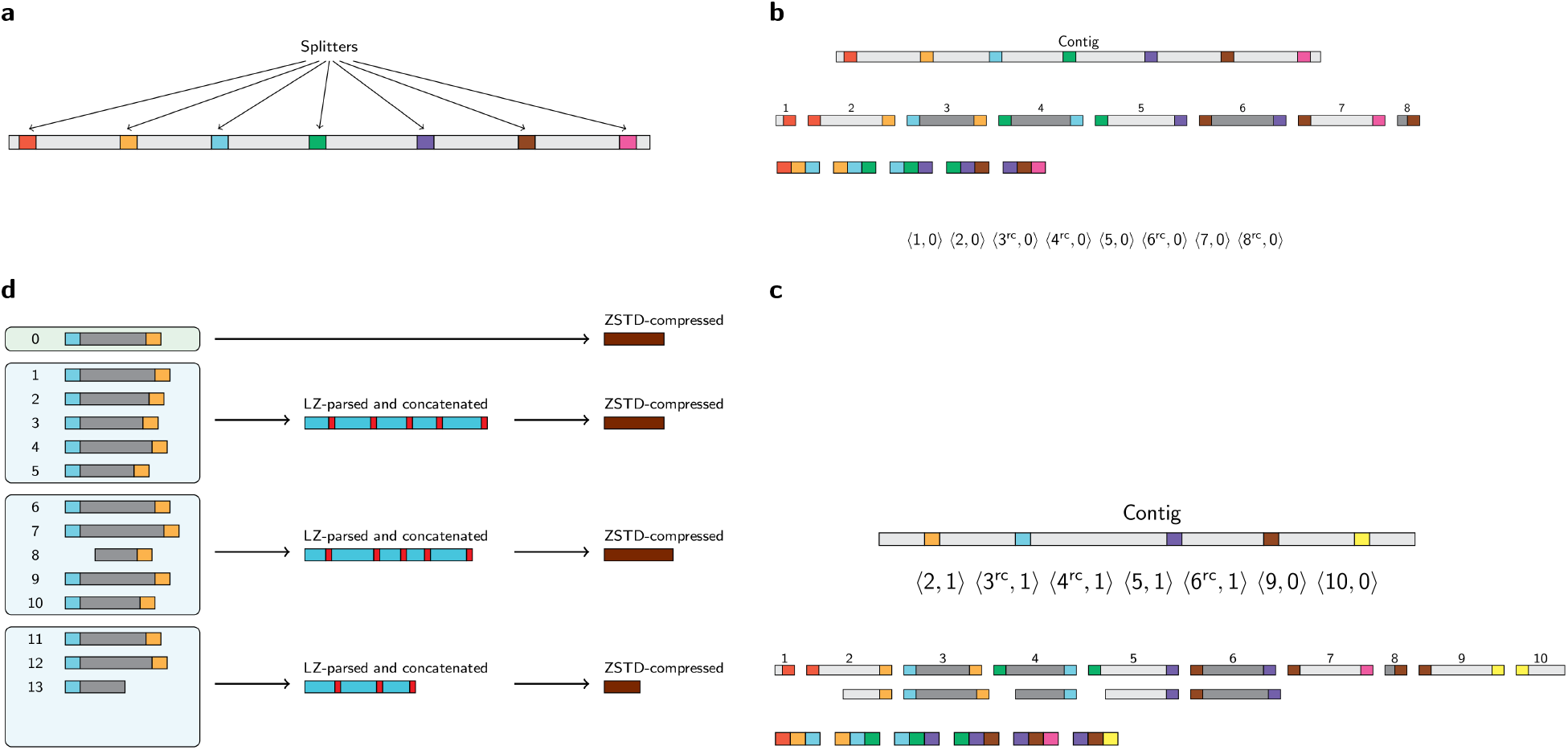
Illustration of the most important stages of the compression algorithm. (**a**) In the first two stages of the compression, we pick some of the *k*-mers of the reference genome (uniformly distributed) as splitters (shown in colors). (**b**) Compression of the first genome. The contigs are split into segments and distributed into groups according to the terminating splitters. For technical reasons some segments are reverse-complemented (dark-gray-marked). The 3-tuples represent the successive splitters in the contigs. (**c**) Compression of remaining genomes. The contigs are split into segments and distributed into groups. If there is no group identified by a pair of terminating splitters (e.g., blue and violet) it is checked if there is a triple of consecutive splitters (blue, *any*, violet). If so, it is assumed that the middle splitter (green) is missing (probably due to some evolutionary event) and (blue, violet) segment is split into two segments: each with only one terminating splitter. Otherwise, e.g., for (brown, yellow) segment, a new group is created. (**d**) Compression of segments in a single group. The reference segment is packed using general-purpose zstd compressor. The remaining segments are LZSS-parsed [13] against the reference segment to find similarities and differences. The short descriptions are concatenated in blocks (red box is a terminator here) and zstd-compressed.

The described strategy is a compromise between compression ratio and access time. It offers very good compression ratios and limits the part of an archive that needs to be decompressed when we want to extract data. Nevertheless, if one is mainly interested in the compression ratio, a bit better strategy could be not using segmentation and allowing the search for similarities in the whole collection, as shown in the MBGC compression for bacterial genomes [6]. However, this would greatly increase the extraction time.

For evaluation, we used several datasets of various species: human, bacterial, and viral (Figure 2e). The largest HPRC dataset consists of 94 human haplotype assemblies, reference genome (GRCh38) and CHM13 genome from the T2T consortium [12]. With each assembly taking approximately 3 GB, the entire dataset requires 293.2 GB space or 79.8 GB when gzipped. AGC compresses it about 200 times to as little as 1.45 GB in about 14 minutes using 32 threads. MBGC (Multiple Bacteria Genome Compressor) [6] is the only tool that can compete in terms of compression ratio. It produces a 1.95 GB archive in longer time (Fig. 2a,c). However, the most important differences between these tools are access to compressed data and the ability to extend the existing archive. AGC is designed to keep the collection in a compact form and extract the samples, contigs or contig fragments when requested, both from command line as well as using programming libraries. For example, the complete human sample can be extracted in less than 3 seconds, independently of the position of the sample in the collection (Fig. 2b). The contigs can be extracted in even a shorter time. This is tens of times faster than MBGC and even faster than extracting from separate gzip archives (Fig. 2b). The existing AGC archive can be easily extended with new samples (Fig. 2d), which takes 10–25 s for a single sample. Experiments with two human datasets, phased (HGSVCp) and unphased (HGSVCu), from the Human Genome Structural Variation Consortium, Phase 2 [5] (36 samples each) lead to similar conclusions. AGC appeared to be quite insensitive to the selection of the reference genome in the human datasets. We examined GRCh38, CHM13 and randomly selected samples as the references and the differences in compression ratio were always less than 7% (Supplementary Table 1).

**Figure 2:**
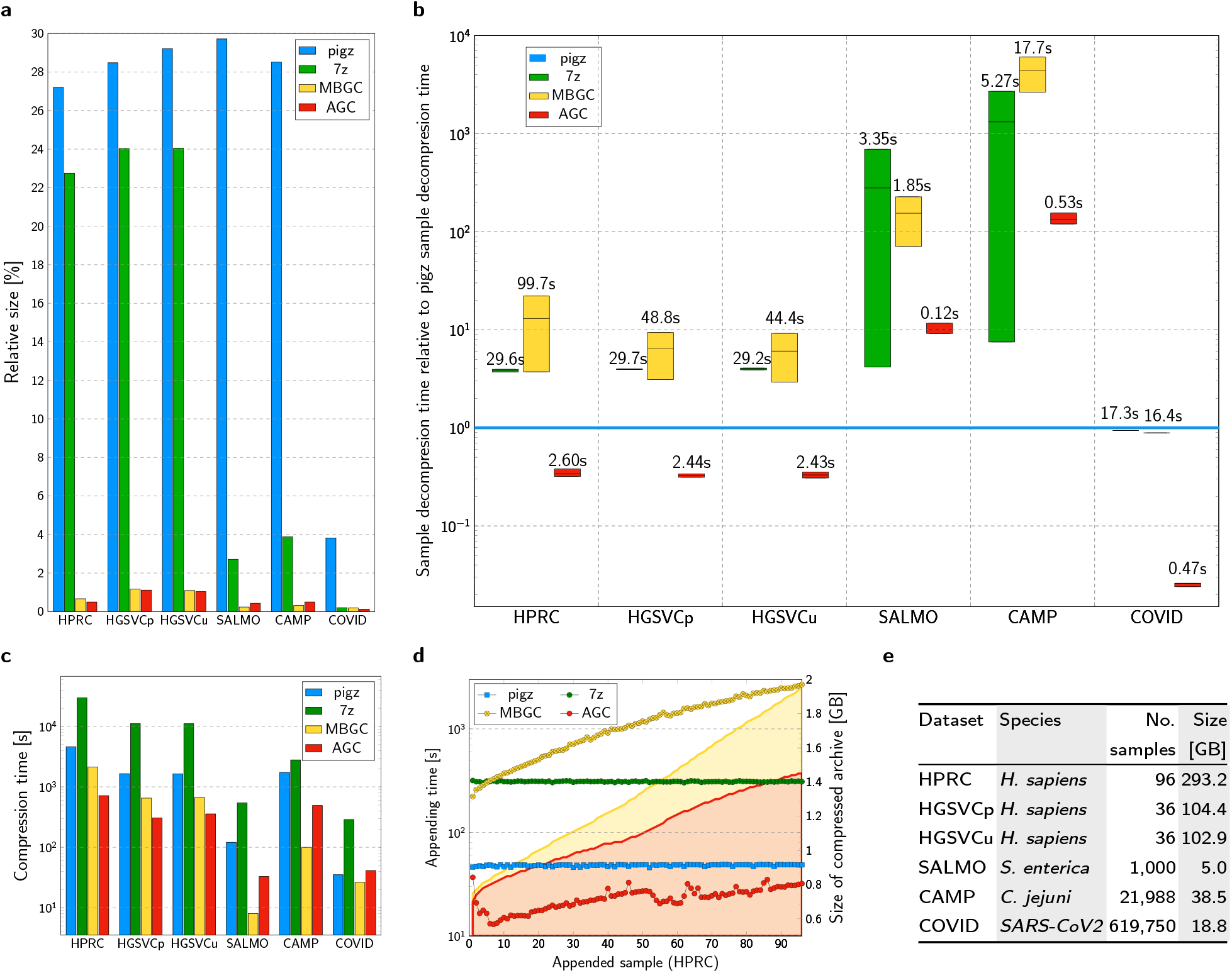
Experimental results. (**a**) Relative sizes of compressed collections of genomes. (**b**) Extraction time of a single sample. The box plots show minimal, median and maximal times relative to decompression time of pigz archives. The labels show median absolute times. (**c**) Compression times of whole collections of genomes. (**d**) Time of appending a single sample to the archive containing given number of HPRC samples (markers). Size of the archives containing given number of samples (filled areas). The filled areas of gzip and 7z are not presented as they are much larger. (**e**) Datasets used in the experiments.

We also experimented with two bacterial datasets, i.e., SALMO (1,000 *Salmonella enterica* samples) and CAMP (21,988 *Campylobacter jejuni* samples) from [3]. Here, MGBC, a tool designed especially for bacterial genomes, wins in terms of compression ratio by producing archives about 40% smaller than AGC. The reason is the high diversity of the bacterial genomes. Therefore, in some contigs there is no splitter, so grouping of segments does not work as well as for human datasets. To partially overcome the problem, for bacterial data we used an adaptive compression mode, in which AGC extends the list of splitters by *k*-mers from contigs without known splitter. Moreover, we used a larger block size (500) and a shorter segment size (1500). However, what is important is that AGC can extract samples in less than a second, while MBGC needs tens of seconds for the larger dataset (Fig. 2b). Furthermore, the extraction time of AGC increases slowly with increasing collection size. This suggests that even for huge collections, AGC can still be seen as a way of on-line access to the samples stored in a very compact form.

In the final test, we evaluated the tools for the highly similar collection of approx. 620k SARS-CoV2 genomes. AGC wins clearly in terms of compression ratio and access time.

Here, we introduce AGC, a versatile package to maintain genome collections in a very compact form. It offers a two-order-of-magnitude reduction of data sizes and allows access to the samples or contigs in seconds or fractions of seconds. Moreover, it allows one to extend the compressed collections by adding new samples. Such a combination of features makes AGC a tool in its own category. This opens new opportunities in the field of (rapidly growing) pangenome projects. AGC archives can be used to distribute pangenome data, store them, and quickly answer queries. Due to the programming libraries provided for popular languages, AGC can be easily integrated with existing pipelines, allowing them to operate on small files.

## Supporting information

Supplementary Worksheet

## Acknowledgements

The work was supported by National Science Centre, Poland, project DEC-2017/25/B/ST6/01525 (SD, AD) and by US National Institutes of Health (grant R01HG010040 and U01HG010961 to HL).

## Author contributions

H.L. defined the problem and requirements of the compression tool for pangenome projects. S.D. designed and implemented the majority of the algorithm. H.L. and A.D. contributed to the design and implementation. H.L., A.D., and S.D. designed the experiments. A.D. performed the experiments and prepared the charts. S.D. wrote the majority of the manuscript. H.L. and A.D. contributed to the manuscript. All authors read and approved the final manuscript.

## Methods

### Algorithm overview

The algorithm is implemented as a multithreading application in the C++17 programming language. It supports all IUPAC codes in the input data (A, C, G, T, U, R, Y, S, W, K, M, B, D, H, V, N). The symbols can be lower- and upper-case but before compression sequences are upper-cased. The application can be run in one of the modes:

- *create*—create an archive from FASTA files,
- *append* —add FASTA files to existing archive,
- *getcol* —extract all samples from archive,
- *getset* —extract a sample from archive,
- *getctg* —extract a contig from archive,
- *listset* —list sample names in archive,
- *listctg*—list sample and contig names in archive,
- *info*—show some statistics of the compressed data.

Initially, the application must be run in *create* mode to build a new archive. The user should provide a collection of genomes to compress and a reference genome. The reference genome can be one of the genomes in the collection or a reference for the species. The user can define the parameters of the archive, for example, *k*-mer length (default: 31), *block size* (default: 50), *segment size* (default: 60,000). Then, the user can extend the archive or ask various types of query.

There are three main stages of compression: (*i*) candidate *k*-mer determination, (*ii*) splitters determination, (*iii*) adding genomes to the archive. The first two occur only in the *create* mode. The last occurs in the *create* and *append* modes.

### Candidate *k*-mers determination

In the first stage of compression, all *k*-mers present in the reference contigs are determined and stored in an array. Then, they are sorted using a fast in-place variant of radix-sort algorithm [8]. Finally, the *k*-mers occurring 2 or more times are removed.

### Splitters determination

Each splitter is a candidate *k*-mer. The number of splitters is more or less the reference genome size divided by *segment size*, so for a human genome and default algorithm parameters it is about 50,000. To determine the splitters, AGC processes contigs of a reference genome one by one from the beginning to the end. For each of them, it looks for the first *k*-mer that is a candidate *k*-mer and stores it as a splitter. Then, it skips *segment size* bases and looks for the next splitter, and so on. The last candidate *k*-mer of a contig is also a splitter.

### Compression of a single genome

The genomes are added to the compressed archive one by one in the same way. The first genome added is, however, the reference genome. The remaining genomes are added in the order provided by the user.

Each genome is compressed contig by contig. At first, each contig is split into *segments*. Segment boundaries are defined by splitters. For technical reasons, a splitter is part of both neighbor segments. There are three types of segments:

- *spt-2* —segment surrounded by two splitters; the majority of segments are of this type,
- *spt-1* —segment with only one splitter; this is usually the case at contig boundaries,
- *spt-0* —segment without any splitter; it can happen that for some short or highly redundant contig it is not possible to localize any candidate *k*-mer; such a segment is always a whole contig. Such a situation can also happen if the whole contig contains a sequence not present in the reference genome.

#### Dealing with *spt-2* segments

Most segments of this type are distributed to groups identified by a pair of splitters they contain. To make this grouping easier, we ‘normalize’ each segment, which means that we compare which of the canonical splitters is lexicographically smaller. Then we reverse-complement the segment if it is necessary to ensure that the smaller splitter is at the beginning of the segment.

The first segment in each group serves as a reference for the group. It is packed: 1, 2, 3, or 4 symbols into a single byte, depending on the alphabet size in the sequence. Then, it is compressed using zstd.

The remaining segments are LZSS-parsed [13] with respect to the reference segment of the group. The (usually) much shorter descriptions of the segments are concatenated into blocks of size *block size*. The blocks are compressed independently using zstd.

However, it is possible that a segment of this type will be split into two segments of type *spt-1*. This could happen if we have a segment with a pair of splitters (*s*_1_, *s*_2_) and there is no group identified by this pair. In this situation, we check if the archive already contains groups identified by splitter pairs (*s*_1_, *s*_3_) and (*s*_3_, *s*_2_) for any splitter *s*_3_. If so, we LZSS-parse the current segment with respect to the reference segments of both groups to find the best division point. Then we split the segment into two *spt-1* segments. If there is no such pair of groups, the current segment starts a new group (especially this is the case when adding the first genome to the archive).

#### Dealing with *spt-1* type segments

We analyze all groups in which at least one splitter is the same as the splitter in the current segment. For each such group, we perform LZSS-parsing to find the group in which the cost of storing the current segment will be the smallest. Then, we add the current segment to this group. A segment of this type can also start a new group if the LZSS-parsing shows that there are no groups similar to the current segment.

#### Dealing with *spt-0* type segments

The segments of type *spt-0* are randomly distributed (but in a deterministic way) into one of 16 groups. Within each group, segments are organized in blocks of size *block size*. Each group is compressed independently using zstd compressor.

### Archive organization

The archive contains compressed blocks from all groups. It also contains descriptions of the contigs, i.e., the ids of groups and within-group ids of segments. These descriptions are also zstd-compressed.

### Extending an archive

The archive can be extended by new samples. In this mode, the archive is partially loaded into the memory. This means that the last blocks of each group are loaded (but not decompressed until it is necessary to add anything to them). We also load the descriptions of the contigs. Then, we proceed as usual when we add new genomes.

### Adaptive mode

For highly divergent species, such as bacteria, better results are possible if the splitters are determined not only in the reference genome but also in the remaining genomes. The processing of the reference is the same as in the default mode. However, when a new genome is added, the *spt-0* type segments are collected in some buffer (not directly stored in the archive). Then, after processing all genome contigs, the *k*-mers in the buffered contigs are counted, duplicates are removed, and also the *k*-mers present in the reference genome are removed to get *sample-candidate k*-mers. Then, these *k*-mers are used to determine new splitters that extend the *global* splitters. After that, the buffered contigs are processed one more time to determine *spt-2, spt-1*, and *spt-0* segments that are handled as usual.

### Decompression of a contig or a sample

To decompress a contig it is necessary to read its description to find which groups will be required. Then we zstd decompress the reference segments of the selected groups. If necessary, we also decompress one of the blocks in a group, get LZSS-parsing of the requested segment, and reconstruct it. In the query, it is possible to restrict to only some part of a contig. In this case, we decompress only the segments necessary to reconstruct the requested part.

Decompression of a sample is just decompression of the contigs it is composed of.

### Block size choice

The block size has a significant impact on the size of the archive, as well as compression and extraction time. We performed a few experiments on the HPRC dataset to measure it. Supplementary Figure 1 shows time necessary to add a single sample to the archive. For example, for *block size* 10 it is usually below 15 s per sample. For larger values, it grows to 25 s but should stay at this level even for larger collections, as this time is dependent mostly on the *block size* and rather little on the collection size. In Supplementary Figure 2, we present how the archive size grows when adding samples one by one for various block sizes. Supplementary Figure 3 shows how long it takes to extract a single sample from an archive for various block sizes. The default *block size* value (50) is a compromise between access time and compression ratio, which should be used for large genomes, such as humans. For bacterial and viral datasets, larger values (e.g., 500) should be a better choice. Finally, Supplementary Figure 4 compares the extraction time from HPRC dataset for the tools examined.

## Data availability

All datasets used in the experiments are publicly available. The details on how to download them are given in the Supplementary Data.

## Code availability

The source code of AGC is available at https://github.com/refresh-bio/agc. The package can be installed via Bioconda at https://anaconda.org/bioconda/agc.

