## Supplementary Data for "AGC: Compact representation of assembled genomes"

### AGC: Compact representation of assembled genomes — supplementary information

April 7, 2022

**3 Environment** **5**

**4 Supplementary Results** **6**

**5 Additional Results** **9**

### 1 Datasets

We used six datasets in experiments. Three sets contain human genome assemblies (HPRC, HGSVCp, HGSVCu), two sets contain bacterial genome assemblies (SALMO and CAMP), and one set contains *SARS-CoV-2* genomes (COVID). Each human and bacterial genome assembly is a separate multi-FASTA file, while all *SARS-CoV-2* genomes are in a single multi-FASTA file (one sequence is one genome).

The Supplementary Worksheet contains exact names and characteristics (size, number of sequences) of all multi-FASTA files used in experiments.

#### 1.1 HPRC

Set contains 96 human genome haploid assemblies, including: CHM13 v.1.1 assembly, GRCh38, and 94 haploid human assemblies released by the Human Pangenome Reference Consortium (HPRC) in 2021 [Nurk et al., 2022].

Data source:
`https://github.com/human-pangenomics/HPP_Year1_Assemblies`

#### 1.2 HGSVCu

Set contains 36 unphased haploid human genome assemblies generated by the Human Genome Structural Variation Consortium (HGSVC), phase 2 [Ebert et al., 2021].

Data source:
`http://ftp.1000genomes.ebi.ac.uk/vol1/ftp/data_collections/HGSVC2/release/v1.0/assemblies/20200612_HHU_assembly-results_CLR_v12/assemblies/unphased/`

#### 1.3 HGSVCp

Set contains 36 phased haploid human genome assemblies generated by the Human Genome Structural Variation Consortium (HGSVC), phase 2 [Ebert et al., 2021]. All sequences were upper-cased before the experiments.

Data source:
`http://ftp.1000genomes.ebi.ac.uk/vol1/ftp/data_collections/HGSVC2/release/v1.0/assemblies/20200612_HHU_assembly-results_CLR_v12/assemblies/phased/`

#### 1.4 SALMO

Set contains 1000 *Salmonella enterica* assemblies chosen from the 661K dataset assembled and characterized in [Blackwell et al., 2021].

Data source: `ftp.ebi.ac.uk/pub/databases/ENA2018-bacteria-661k`

#### 1.5 CAMP

Set contains 22,988 *Campylobacter jejuni* assemblies chosen from the 661K dataset assembled and characterized in [Blackwell et al., 2021].

Data source: `ftp.ebi.ac.uk/pub/databases/ENA2018-bacteria-661k`

#### 1.6 COVID

Set contains 619,750 complete *SARS-CoV-2* genomes downloaded from the National Center for Biotechnology Information (NCBI) at the end of 2021.

Data source:
`https://www.ncbi.nlm.nih.gov/datasets/coronavirus/genomes/`
Data availability:
`https://zenodo.org/record/5826274/files/sars-cov-2_ncbi-620k.fa.xz?download=1`

### 2 Examined programs

The following programs were used in the experimental part. Running parameters are also given.

#### 2.1 pigz v. 2.6

- Compression

```
pigz -9 -p 32 [file]
```

- Decompression

```
pigz -9 -d [archive]
```

## 2.2 7z v. 16.02

In case of HPRC, HGSVCu, HGSVCp and COVID datasets each multi-FASTA file was compressed and decompressed separately, as compressing files from single dataset together did not reduce the total archive size, while increasing the access time (and COVID dataset is a single file):

- Compression

```
7za a -mx9 -mmt32 [archive] [file] > /dev/null
```

- Decompression

```
7za e  [archive] > /dev/null
```

In case of SALMO and CAMP dataset all multi-FASTA files from the same dataset were compressed together:

- Compression

```
7za a -mx9 -mmt32 [archive] @[files_list]  > /dev/null
```

- Decompression (whole set)

```
7za e   [archive] > /dev/null
```

- Decompression (single file)

```
7za e  [archive] [file]  > /dev/null
```

#### 2.3 MBGC v. 1.2.2PE

There are three possible compression modes for MGBC [Grabowski & Kowalski, 2022] (speed: 0; default: 1; repo: 2; max: 3), but as the "speed" and the "default" modes did not work for human data and the preliminary test showed that the "max" mode is significantly slower in sample decompression than the "repo" mode, we used the "repo" mode in all experiments described in this paper.

In the case of HPRC, HGSVCu, HGSVCp, SALMO and CAMP datasets multiple file compression was used:

- Compression (whole set)

```
mbgc  -c 2 -t 32 [files_list] [archive] > /dev/null
```

- Decompression (whole set)

```
mbgc  -d -t 32 -l [line_length] [archive] [output_folder] > /dev/null
```

- Decompression (single sample / file)

```
mbgc  -d -t 32 -f [sample] -l [line_length] [archive] [output_folder] > /dev/null
```

- Appending a sample / file

  There are no options for archive extension. To add a sample to an existing MGBC archive, it is necessary to decompress the archive and compress the extended set of files.

In case of COVID dataset, single-file compression was used:

- Compression

  ```
  mbgc -c 2 -t 32 -i [file] [archive] > /dev/null
  ```

- Decompression

  ```
  mbgc  -d -t 32 -l [line_length] [archive] [output_folder] > /dev/null
  ```

##### 2.4 AGC v 2.0

For all datasets the first sequence in the set was used as a reference.

In case of HPRC, HGSVCu and HGSVCp datasets multiple file compression was used with the default parameters (batch size $b = 50$, segment size $s = 60000$):

- Compression

  ```
  agc create -t 32 -i [files_list] [reference] > [archive]
  ```

- Appending a sample / file

  ```
  agc append -t 32 -o [new_archive] [archive] [file]
  ```

In case of SALMO and CAMP datasets multiple file compression was run in adaptive mode with batch size $b = 500$ and segment size $s = 1500$:

- Compression

  ```
  agc create -t 32 -a -b 500 -s 1500  -i [files_list] [reference] > [archive]
  ```

- Appending a sample / file

  ```
  agc append -t 32 -a -o [new_archive] [archive] [file]
  ```

In case of COVID dataset, as all samples are concatenated in a single file, a dedicated '-c' option was used for compression:

- Compression

  ```
  agc create -s 3000 -b 10000 -t 32 -c [refrerence] [file]
  ```

Decompression for all datasets:

- Decompression (whole set)

  ```
  agc getcol -t 32 -l [line_len] -o  [out_folder] [archive]
  ```

- Decompression (single sample / file)

  ```
  agc getset -t 32 -l [line_len] [archive] [sample] > [output_file]
  ```

### 3 Environment

The machine used in the tests was of the following configuration:

- AMD Ryzen Threadripper 3990X 64-Core Processor, 128-Threads, 2.9GHz
- 256 GiB RAM,
- 3.6 TB NVMe disk

For compilation, we used G++ v. 11.2.0. The machine was running Debian 12. For some tests (i.e. sample decompression times) we used a temporary file storage facility (tmpfs) of size 128 GB.

### 4 Supplementary Results

Table 1: Archive sizes for human datasets for reference genome randomly selected from the collection and remaining genomes appended in a random order.

| Reference sequence | Archive size [MB] |
|---|---|
| **HPRC dataset** | |
| chm13.draft_v1.1.fasta | 1454 |
| GCA_000001405.15_GRCh38_no_alt_analysis_set.fna | 1464 |
| HG00621.paternal.f1_assembly_v2_genbank.fa | 1523 |
| HG01071.paternal.f1_assembly_v2_genbank.fa | 1473 |
| HG01243.paternal.f1_assembly_v2_genbank.fa | 1544 |
| HG02109.paternal.f1_assembly_v2_genbank.fa | 1493 |
| HG03486.paternal.f1_assembly_v2_genbank.fa | 1496 |
| **HGSVCp dataset** | |
| v12_HG00096_hgsvc_pbsq2-clr_1000-flye.h1-un.arrow-p1.fa | 1162 |
| v12_HG00731_hgsvc_pbsq2-clr_1000-flye.h2-un.arrow-p1.fa | 1150 |
| v12_HG01596_hgsvc_pbsq2-clr_1000-flye.h2-un.arrow-p1.fa | 1170 |
| v12_NA19239_hgsvc_pbsq2-clr_1000-flye.h2-un.arrow-p1.fa | 1164 |
| v12_NA20847_hgsvc_pbsq2-clr_1000-flye.h1-un.arrow-p1.fa | 1179 |
| **HGSVCu dataset** | |
| v12_HG00096_hgsvc_pbsq2-clr_1000_nhr-jax27.fasta | 1066 |
| v12_HG00731_hgsvc_pbsq2-clr_1000_scV12-hhu26.fasta | 1101 |
| v12_HG01596_hgsvc_pbsq2-clr_1000_scV12-hhu27.fasta | 1170 |
| v12_NA19239_hgsvc_pbsq2-clr_1000_scV12-hhu26.fasta | 1116 |
| v12_NA20847_hgsvc_pbsq2-clr_1000_nhr-hhu26.fasta | 1063 |

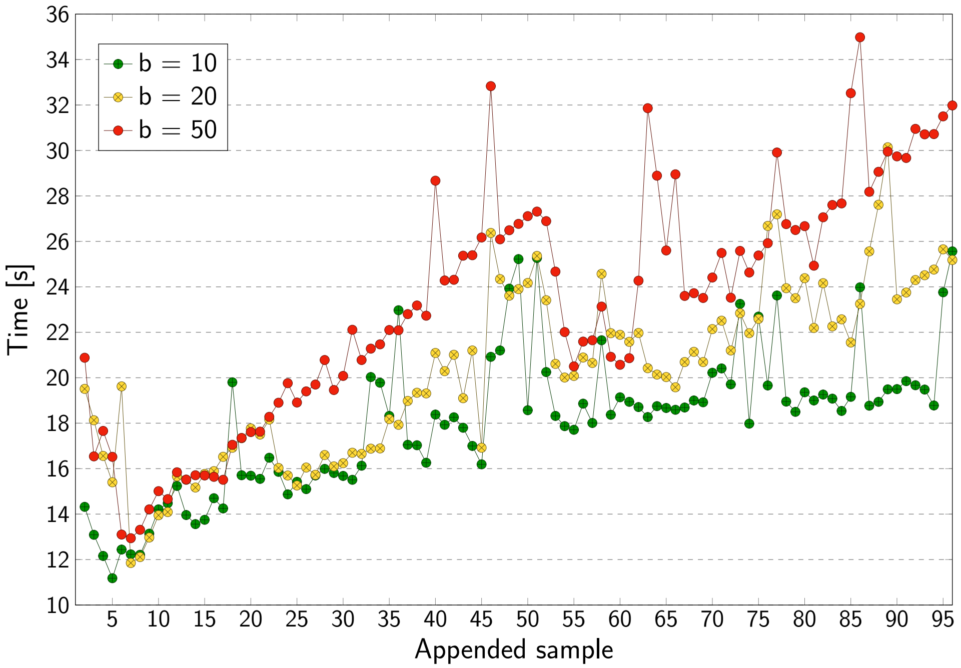

Figure 1: Time of extending the collection by adding a single HPRC human sample at a time. The series show impact of block size.

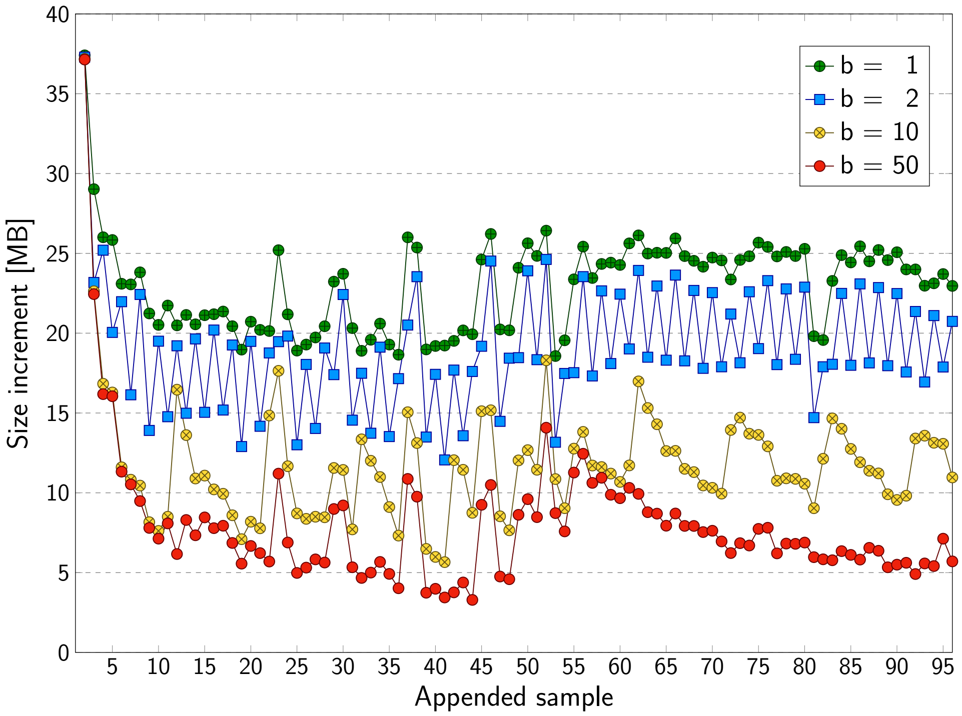

Figure 2: Archive space increase after extending the collection by adding a single HPRC sample at a time. The series show impact of block size.

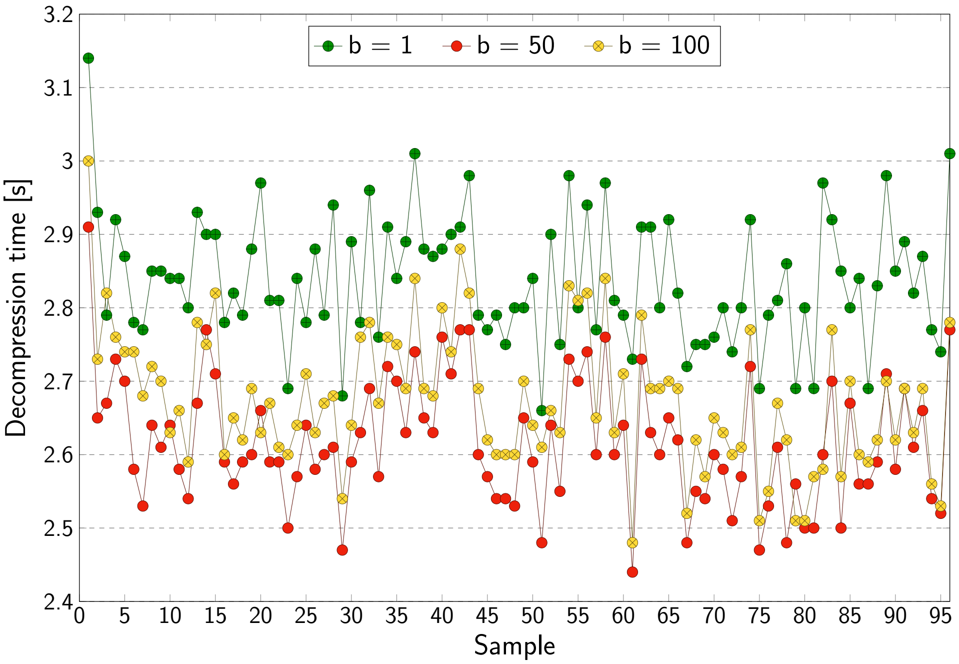

Figure 3: Decompression time of a single HPRC sample. The series show impact of block size.

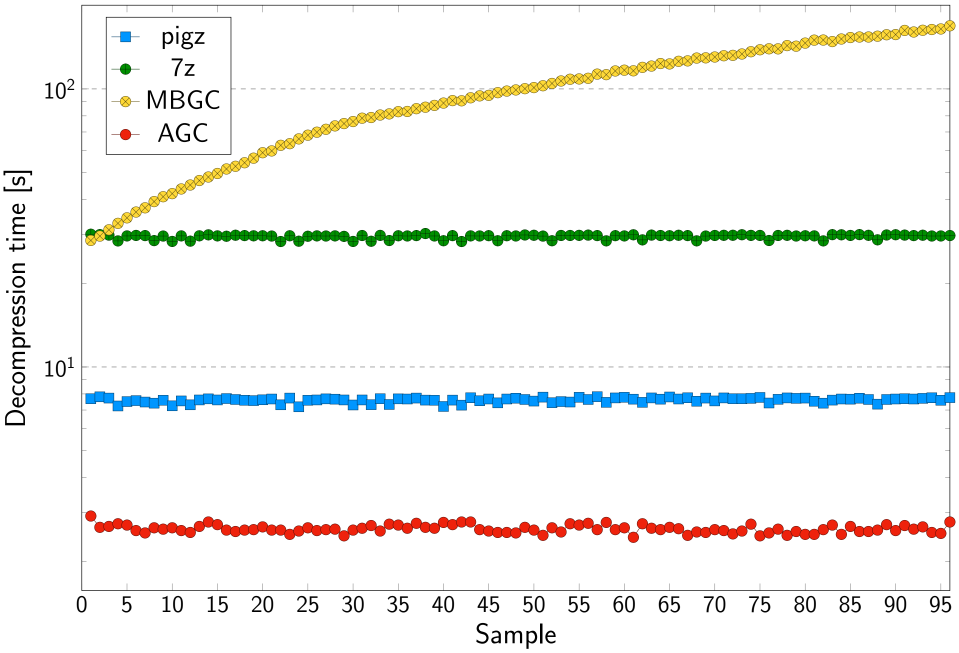

Figure 4: Extraction time of a single HPRC sample as a function of sample position in the archive.

#### 5 Additional Results

Additional results are given in the Supplementary Worksheet.
